# Systematic inference of regulation by protein kinases finds surprising level of transcription factor deactivation

**DOI:** 10.1101/2021.09.03.458813

**Authors:** Christian Degnbol Madsen, Jotun Hein, Christopher T. Workman

**Affiliations:** Department of Biotechnology and Biomedicine, Technical University of Denmark, 2800 Kongens Lyngby, Denmark; Department of Statistics, University of Oxford, Oxford OX1, UK

## Abstract

Gene expression is controlled by pathways of regulatory factors often involving the activity of protein kinases on transcription factor proteins. Despite this well established mechanism, the number of well described pathways that include the regulatory role of protein kinases on transcription factors is surprisingly scarce in eukaryotes.

To address this, *PhosTF* was developed to infer functional regulatory interactions and pathways in both simulated and real biological networks, based on linear cyclic causal models with latent variables. *GeneNetWeaverPhos*, an extension of *GeneNetWeaver*, was developed to allow the simulation of perturbations in known networks that included the activity of protein kinases and phosphatases on gene regulation. Over 2000 genome-wide gene expression profiles, where the loss or gain of regulatory genes could be observed to perturb gene regulation, were then used to infer the existence of regulatory interactions, and their mode of regulation in the budding yeast *Saccharomyces cerevisiae*.

Despite the additional complexity, our inference performed comparably to the best methods that inferred transcription factor regulation assessed in the *DREAM4* challenge on similar simulated networks. Inference on integrated genome-scale data sets for yeast identified ∼8800 protein kinase/phosphatase-transcription factor interactions and ∼6500 interactions among protein kinases and/or phosphatases. Both types of regulatory predictions captured statistically significant numbers of known interactions of their type. Surprisingly, kinases and phosphatases regulated transcription factors by a negative mode or regulation (deactivation) in over 70% of the predictions.

**Author summary:** In this work we addressed the challenging problem of inferring regulation by protein kinases and phosphatases via their activity on transcription factors. Although many protein kinase activity predictors have been developed for classes of protein kinases on specific amino acids within target sequences, our approach (PhosTF) provides predictions of regulatory activity for specific protein kinases and phosphatases on specific transcription factors. Our inference approach achieves this using the functional output observed in gene expression data of gene knock out stains, along with known transcription factor regulatory interactions. We formulated and tested a model for inference of regulation as well as a model for simulation of genes expression, transcription and translation. The simulation was used for in-silico validation of the inference method, which performed comparably to the best performers on simpler inference problem in the DREAM4 competition. The inference method was then applied to yeast expression data, with significant validation by known kinase/phosphatase interactions. Over 15300 novel regulatory interactions were predicted, suggesting that kinase activity provided a surprising level of repression of gene expression, either through the deactivation of activators or the activation of repressors.

## Introduction

Gene regulation is central to a cell’s ability to respond and adapt to changes in its environment. The control of transcription rates are directly regulated by transcription factors (TFs), and indirectly by chromatin state, cell signalling and other regulatory factors. Modulation of TF activity is often achieved through phosphorylation or dephosphorylation by protein kinases (PKs) or phosphatases (PPs), and TFs represent one of the most phosphorylated classes of proteins [1]. Regulation by TFs can be mapped from protein-DNA binding experiments, e.g. by chromatin immunoprecipitation (ChIP) based methods, while protein kinase and phosphatase (KP) regulation can be inferred from protein-protein binding as measured by yeast two-hybrid, or co-immunoprecipitation and mass spectrometry-based methods. These technologies suffer from false negatives due to the transient nature by which kinases and phosphatases bind their targets, as well as false positives [2]. Online databases containing protein interactions will sometimes report whether the data is collected from low- or high-throughput experiments, or whether they were observed reproducibly in multiple experiments, but information about data quality or functionality is often limited [3]. To infer functional regulatory interactions, one can draw from multiple sources of data, both protein binding data and evidence of regulation from mRNA transcript levels. In particular, when comparing the transcript levels from mutant strains, e.g. gene knock-out or overexpression strains, to their background strains, the output of regulatory pathways can be observed by the resulting changes in mRNA levels. The loss or gain of a TF or KP gene will often generate altered transcript levels that imply functional regulation or a regulatory dependency between the perturbed regulator and the gene with an altered mRNA level [4]. Inference of functional regulation can be achieved through the use of these diverse data types as evidenced by a number of approaches used in the DREAM4 challenge [5].

To this aim, methods for the inference of TF-based regulatory networks were evaluated in the DREAM4 challenge [5]. To do this, mRNA concentrations were simulated with the software GeneNetWeaver from a known network using differential equations describing mRNA and protein concentration gradients [6]. In this way, all regulators can be deleted or overexpressed (in silico) in turn and new steady-state mRNA output can be generated for each. However, GeneNetWeaver does not take phosphorylation or other post-translational modification into account. The focus of our approach was to extend the inference of TF-based regulatory networks to include the activity of kinases and phosphatases, and to apply this method to the model budding yeast *Saccharomyces cerevisiae*, which has been thoroughly mapped for protein interactions.

The majority of efforts to infer regulatory networks have focused on TFs binding to promoter regions of their target genes. These networks are often modeled as directed acyclic graphs (DAG) of TF nodes interacting with nodes representing target genes. Applied in this biological context, each node value represents a protein’s concentration and each edge the direct regulatory effect, or activity, from node to node. Modelling regulation as a DAG has limitations on the extent to which graph edges represent causality and physical interaction, since gene regulation is highly cyclic as target gene products are often regulators themselves. The *linear cyclic causal models with latent variables* (LLC) approach was specifically designed to address inference of causality in cyclic graphs and has been applied to infer TF regulatory networks [7].

Methods have also been proposed that combine multiple likelihood functions for numerous types of evidence [3]. In such cases, maximum likelihood ratios can be calculated for each potential regulatory interaction (edge) by describing the likelihood ratios through factor graphs. Inferring regulation from KPs is much more challenging since they do not regulate mRNA production rates directly, but rather modulate protein activity of other potential regulators. Although there have been many kinase prediction approaches (NetPhos, NetPhospan, PhosphoPredict) most have focused on the phosphorylation site prediction often without the ability to identify which kinase was likely responsible for the phosphorylation. Recent studies utilizing mass spectrometry based proteomics or phosphoproteomics have investigated the activity of KPs in knockout studies [8] [9]. Though these studies endeavour to infer KP activity, they do so relative to the effects on metabolic processes induced by environmental perturbations, thus associating KPs to cellular responses rather than specific regulators. Approaches to infer regulation between human protein kinase (PK-substrate regulatory interactions) has revealed extensive circuits of kinases [10]. Their approach was based on an ensemble method combining multiple kinase-substrate scores and included phosphosite data for specific kinases or kinase families measured by phosphoproteomics, and gene expression data for quantifying co-expression and co-regulation. The combination of multiple data sources was leveraged to achieve high performance, but limits the method to organisms where extensive kinase-phosphosite data is available.

In this paper we have developed PhosTF, which builds on the LLC method. PhosTF allows for inference of indirect regulation caused by kinase and phosphatase activity on TFs. Although KP regulation is difficult to infer from any single knockout, their activity can be inferred in combination with TF knockouts as illustrated in Fig 1. We extended GeneNetWeaver simulations in GeneNetWeaverPhos to include regulation by phosphorylation, and describe a new method to infer gene regulatory networks based on a corpus of protein interaction data and a large compendium of knockout and overexpression transcription profiles.

**Fig 1.**
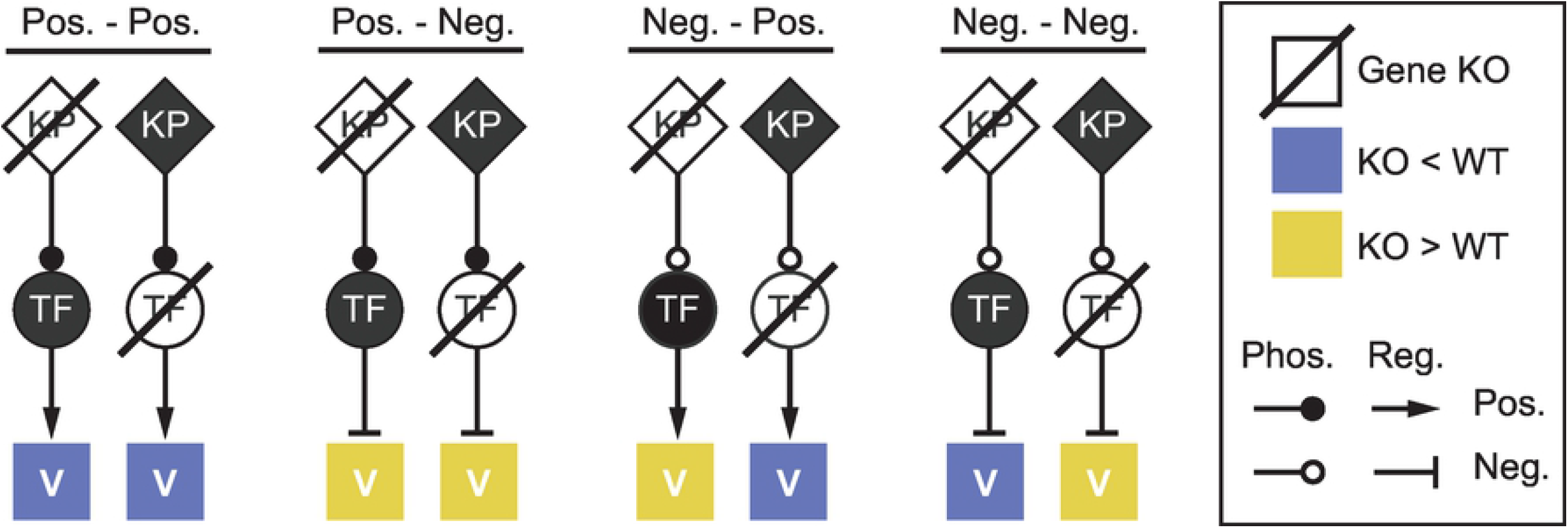
Effects of gene deletion on gene expression. Schematic of gene expression levels for a target gene ‘V’ relative to wildtype when a gene for either a PK or TF is deleted. All combinations of positive (activating) and negative (repressing) regulation (Pos. or Neg. respectively) are shown. Activating or repressing phosphorylation (Phos.) are indicated with closed or open circles and regulation by TFs (Reg.) are indicated with pointed and flat arrowheads.

## Results

The inference method PhosTF was developed and tested on both simulated and experimental data sets. Initially, small-scale simulated data sets were used to illustrate the inference performance, where the regulatory interactions were known. Subsequently, PhosTF was tested on a large compendium of experimental data collected for budding yeast, *S. cerevisiae*.

PhosTF was tested on defined regulatory networks and their simulated output in order to allow for inference where the regulation was known and to allow for the assessment of performance. Given the right parameters, and in particular the appropriate regularization strength, PhosTF was able to correctly infer all edges on the vast majority of the small 5-10 node networks tested. Initially, data was simulated by iterating the PhosTF model equations Eq (3), in which case it was possible to also infer the exact edge weights (within rounding error), which defined the regulatory effects of one gene on another.

### Inference example with ambiguity

In development of the method, extra attention was given to difficult examples, where perfect inference was not always achievable. It was observed for inference on some small networks that if two KPs were similar in their regulatory roles, it could become impossible to distinguish between one regulating the other, or both regulating the same target. The small example shown in Fig 2, with 3 KPs, 2 TFs, 1 target gene and 6 regulatory interactions, illustrates an example where two KPs have a similar role. Despite the ambiguity between KP_1_ and KP_2_, all 6 known regulatory interactions were correctly predicted along with two extraneous edges (false positives). These extra edges give KP_1_ and KP_2_ the same regulatory interactions in the network, both regulating each other and TF_1_.

**Fig 2.**
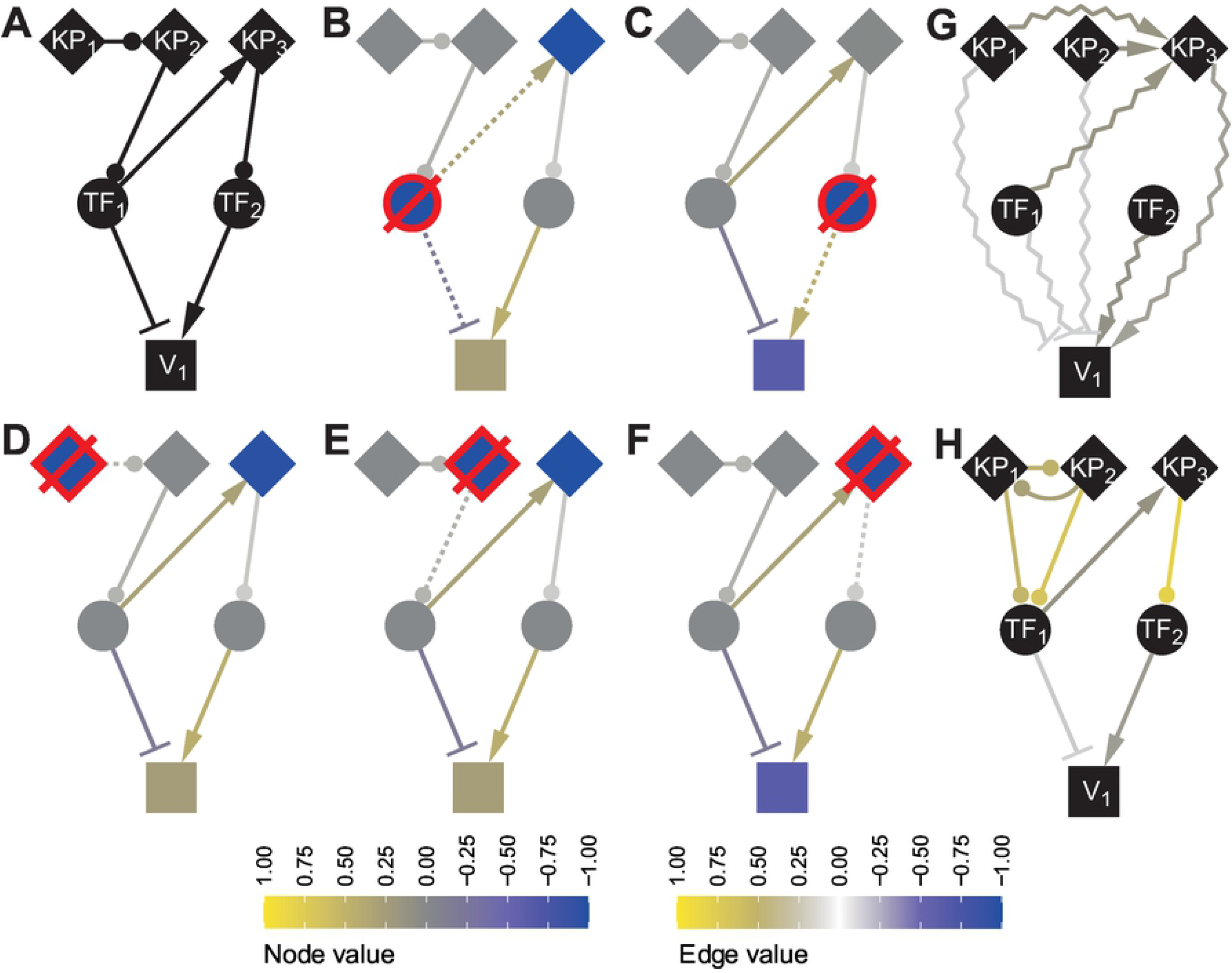
Small example network with unresolvable ambiguity. The true network (A), KPs (diamonds), TFs (circles) and target gene ‘V_1_’. The resulting mRNA outputs from simulated KO experiments are shown in (B-F) and represents the data used for inference. A graph representing the cumulative (total) influence through all pathways is shown in (G), and the inferred regulatory interactions are shown in (H). The node value color scale applies to (B-F) where colors show mRNA log fold-change values for each knockout. The Edge value color scale shows the “true” regulatory weights in (B-F) and the inferred values in (H). The knockout protein is indicated with a red border and strike-through. Dotted edges indicate the direct effects that were removed by the knockout.

The way in which regulation is inferred is illustrated in the total effect graph shown in Fig 2G, which contains the sum of all paths from one node to another [11]. If there is only a single sample of each knockout experiment, then the total effect from node *j* to *i* is simply 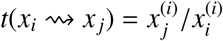 where superscript indicates the knockout. In this case the total effects from node KP_1_ and KP_2_ have become indistinguishable (as seen in Fig 2D,E,G) which makes it difficult to infer the exact regulatory mechanism from perturbation data alone. Due to the ambiguity, multiple potential edges are given non-zero weights. However this behaviour can be modified by changing the regularization of KP edges, or tuning the hyperparameter of the loss function (see Loss in Methods). The log fold-changes shown in Fig 2B-F were calculated from simulated steady-state values (after convergence) of the ODE model defined in Eq (8) for the knockout and wildtype (background) strains.

### Inference performance on simulated data

PhosTF performance was assessed on 25 randomly generated regulatory networks (see Network Construction for Simulation in Methods). The 100-node networks had on average: 20 TFs and 20 KPs, 13 *d*(KP, TF), 13 *d*(KP, KP), 25 *d*(TF, KP), 21 *d*(TF, TF), and 102 *d*(TF, *O*), where *d*(Sources, Targets) denotes directed edges from *v*_*j*_ ∈ Sources to *v*_*i*_ ∈ Targets and *O* is the set of non-regulating nodes.

Results of a Receiver Operator Characteristic (ROC) analysis, Fig 3A, showed that KP regulation could be inferred more accurately for positively regulating edges between KPs, *d*(KP, KP, +), when compared to negative regulation, *d*(KP, KP, −). Unsurprisingly, the KP regulation of TFs was easier to detect than for KPs on other KPs, since the latter are more distant in a regulatory pathway from the transcriptionally regulated genes. Again, differences in prediction performance were observed between positive and negative modes of KP regulation of transcription factors. Surprisingly high performances were observed for all types of *d*(KP, TF) regulation. When selecting edges by the amplitude of the weights (| *w*_*i j*_|) an area under the ROC curve (AUC) of 0.89 was achieved. Even higher performances were observed for positive (*d*(KP, TF, +), AUC=0.94) and negative regulation (*d*(KP, TF, −), AUC=0.90) when assessed separately. We observed similar prediction performance for transcriptional regulation (*d*(TF, *V*), AUC=0.84) compared to the methods assessed in the DREAM4 challenge (TF regulation only), where LLC had an overall AUC=0.76 [11], and an AUC=0.83 was achieved for the best performance on the original 100-node networks by team “ALF” [12].

**Fig 3.**
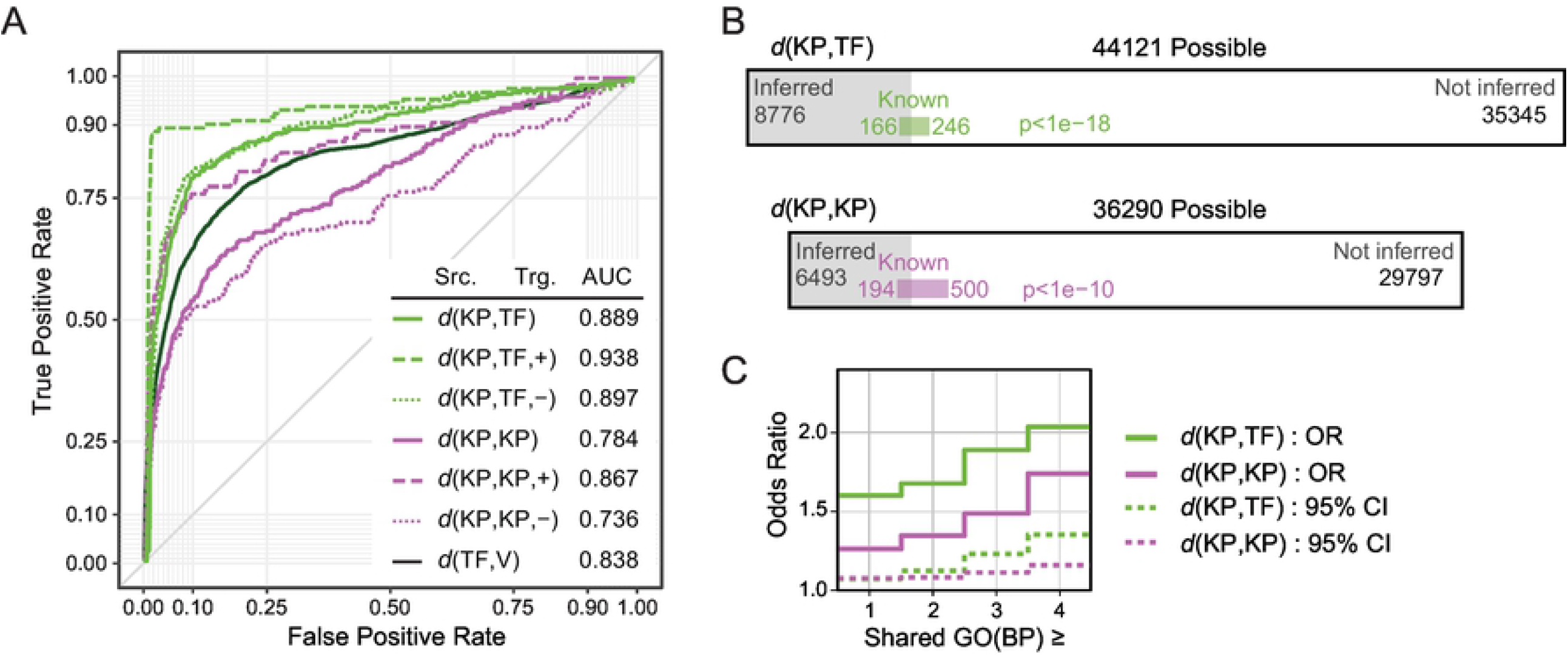
Performance evaluations. Edge inference assessment on simulated networks (A) with complete knowledge of “true” interactions, and on yeast (B and C) where very limited examples were known. ROC curves illustrating edge inference performance pooled from all 25 simulated networks, each containing 100 nodes. Different types of regulation were assessed independently as indicated in the legend. Gene set ‘V’ refers to any gene. Area proportional diagrams for yeast inference (B). Performance on evaluation set of edges visualized for ∼80 substrates/KP. Fisher’s exact test *p*-values represent the chance of observing the performance by random chance. Odds ratio for the number of shared GO Biological Process (BP) terms between interacting genes (C) based on the number of shared GO terms for inferred compared to uninferred edges. Dashed lines show the 95% confidence intervals of the odds ratios.

### Inference performance on yeast data

Regulatory network inference for yeast was applied to a large curated set of yeast gene knockout and overexpression studies (transcriptomics), direct measurements of phosphorylation sites, and binding of transcription factors to DNA (see Yeast Data in Methods). While the regulation from transcription factors is well studied in *S. cerevisiae* (>20,000 interactions), the number of known protein kinase and phosphatase (KP) regulatory interactions on transcription factors, or between KP proteins is very small (412 and 694 interactions respectively).

In an effort to focus on the inference of KP regulation, regulatory networks were inferred for KP interactions only. For this reason, only weights for the known TF regulatory interactions were estimated, thus reducing the complexity of the inference problem. By contrast, all possible *d*(KP, TF) and *d*(KP, KP) edges were included in the inference. Edge weights were trained (see Network Construction in Methods) and then separately filtered for *d*(KP, TF) and *d*(KP, KP) edges with a false discovery rate (FDR) threshold *q* < 0.05 resulting in ∼80 substrates per KP, which is comparable to the average number of kinase targets previously reported (47) [1]. In total, 8776 *d*(KP, TF) and 6493 *d*(KP, KP) were predicted at this FDR threshold for 146 protein kinases and 45 protein phosphatases.

The two types of inferred KP edges were evaluated relative to the limited set of known interactions (Fig 3B) using Fisher’s exact test. For both *d*(KP, TF) and *d*(KP, KP) edges, the predicted set of interactions was significantly enriched (*p* < 10^−10^) for known interactions. These results were compared to predictions made by the protein sequence based substrate prediction method NetPhorest [13]. NetPhorest included 33 protein kinases which could be found in the evaluation set (where the source node is among the 33). The top scoring NetPhorest edges were selected in a number proportionally to the number of inferred edges shown in Fig 3B and resulted in Fisher’s exact test *p* < 0.05 for both *d*(KP, TF) and *d*(KP, KP) NetPhorest predictions. The NetPhorest predictions contained 30% of the possible known edges compared to the 40% captured by PhosTF.

The regulatory interactions inferred by PhosTF were also evaluated relative to shared Gene Ontology terms of regulator-target pairs. The odds ratio of having one or more shared GO slim Biological Process terms (GO(BP)) for the source and target genes of an inferred edge (compared to an uninferred edge) were 1.52 and 1.20 for *d*(KP, TF) and *d*(KP, KP) respectively, see Fig 3C. This odds ratio was observed to increase with the minimum number of shared GO terms. For example, the odds ratio increased to 2.05 for *d*(KP, TF) and 1.67 for *d*(KP, KP) edges when source and target genes shared 4 or more GO terms. All odds ratio estimates were outside the standard 95% confidence interval (CI), which implied a biological significance to both types of predictions with respect to capturing regulatory relationships between proteins functioning together in known biological pathways.

Due to the higher prediction performance of *d*(KP, TF) edges relative to *d*(KP, KP) edges, further analyses were performed for predicted regulatory interactions between protein kinases/phosphatases and their targeted transcription factors. Counts for protein kinase (PK) and phosphatase (PP) edges were summarized in Fig 4A reflecting the combinations in Fig 1 for the positive or negative regulation of either a positively or negatively regulated TF-regulon. This simplified the role of a TF to have a single mode of regulation so as to focus on the role of KP regulation and avoid counting all combinations of paths from the TF. The proportions of these predicted regulatory interactions on TFs was tested relative to the expected counts in two χ^2^ tests (separately for *d*(PK, TF) and *d*(PP, TF)). The expected counts were calculated under the null hypothesis where KP and TF modes of regulation were independent. For instance, the expected number of positive *d*(PK, TF) edges onto a TF activator (Pos.-Pos.) is the fraction of *d*(PK, TF) edges that are positive × the fraction of activating TFs, scaled by the total number of *d*(PK, TF) edges. It was found that negative PK-regulation of TFs that negatively regulate gene expression of their regulons (Neg.-Neg.) were over-represented by 21% (*p* < 1 × 10^−7^). The increased number of Neg.-Neg. pathways is contrasted with a relative under-representation of Pos.-Pos. pathways which were found to be 10% less than expected. The observed distribution of *d*(PP, TF) edges was not observed to differ from the expectation based on a similar calculation.

**Fig 4.**
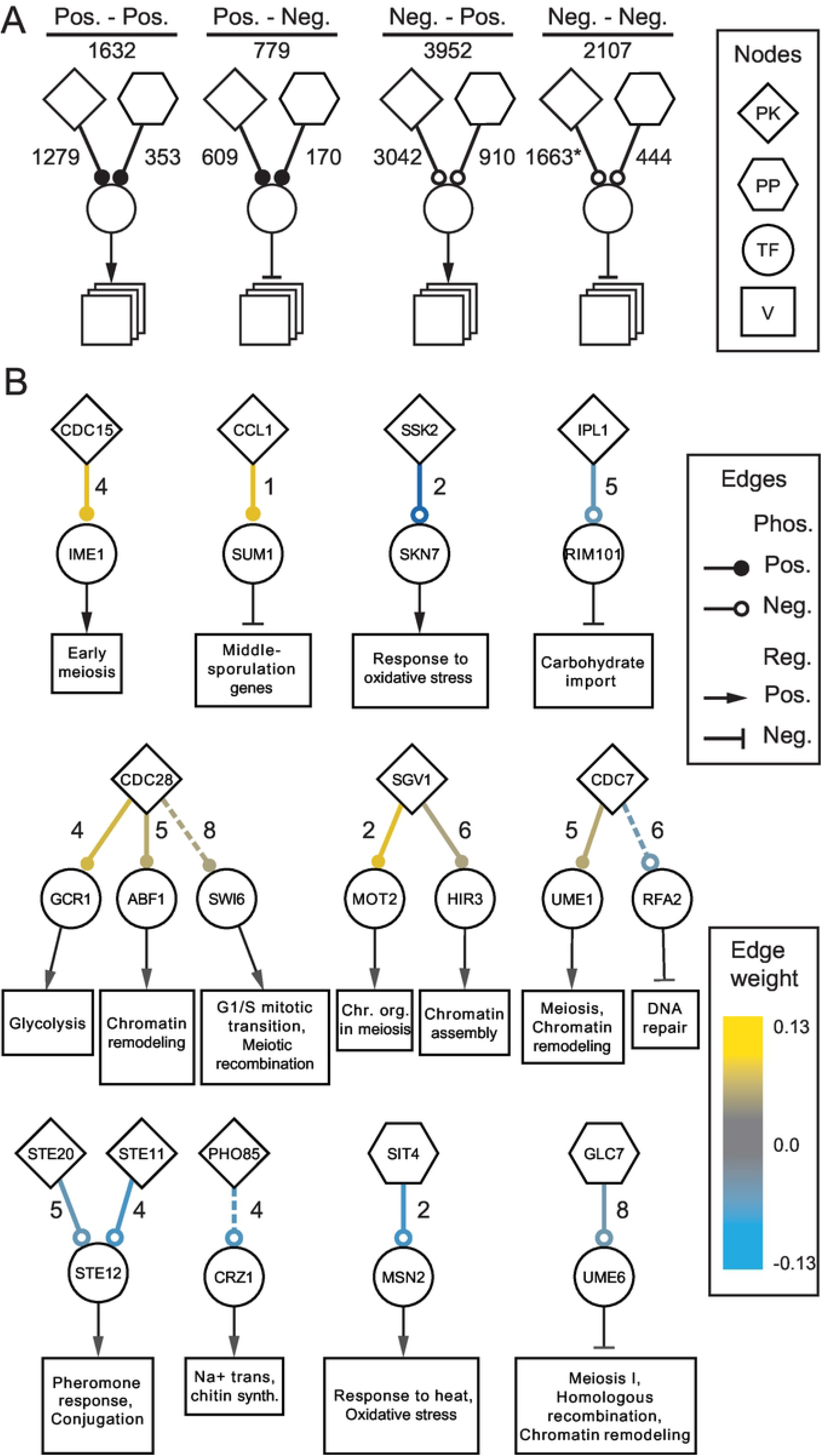
Inferred regulatory pathways. A summary of *d*(KP, TF) edges is shown in (A). Counts of inferred edges for each combination of regulation mode for *d*(PK, TF) or *d*(PP, TF) to either a transcriptional activator or repressor. Counts statistically larger than expected are marked with asterisk. The top scoring *d*(KP, TF) edges with shared OG terms are shown in (B). The number of GO terms shared between KP and TF shown next to each edge. Candidate edges have large absolute weights selected using various thresholds for the number of shared GO terms. Dotted lines indicates edges present in the evaluation set, i.e. known regulatory interactions (true positives). Boxes represent regulons for the TFs and are labeled with representative significant biological process GO terms or the process the TF is known to regulate. The size of the box represents the number of genes in the regulon.

The top positively and negatively regulated KP pathways with shared GO terms were selected as candidates for further investigation and are shown in Fig 4B (see Methods). Of the resulting 16 *d*(KP, TF), three were known (in the validation set) and shared at least four GO(BP) terms. Of the remaining 13, five were found to be directly associated in the STRING database (scores > 0.5), albeit only through types of evidence other than physical interactions. This left only eight interactions (50%) which appear to be novel observations.

## Methods

### PhosTF model definition

The inference is centered around a linear model of the influence nodes have on the values of other nodes, and how interventions on this graph can be used to infer the presence of edges in the graph, as well as their mode of regulation (positive or negative).

A number of node (vertex) sets were defined that represent the potential regulatory role of a gene, or its ability to be modeled by this regulation (Table 1).

**Table 1.** Node set definitions.

| Set | Description |
| --- | --- |
| TF | all transcription factor genes |
| PK | all protein kinase genes |
| PP | all protein phosphatase genes |
| KP | $\text{PK} \cup \text{PP}$ |
| KPTF | $\text{KP} \cup \text{TF}$ |
| $O$ | genes with no regulatory role |
| $V$ | all genes in the model. $\text{KP} \cup \text{TF} \cup O$ |

### Equilibrium equations

The following are the difference equations used to model the node attributes as a function of discreet time steps.

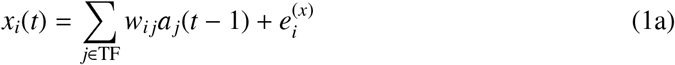

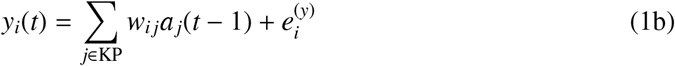

*x*_*i*_(*t*) represent the relative mRNA concentrations, specifically log_2_ fold-change mRNA concentration for a mutant relative to wildtype. Equivalently, *y*_*i*_ represents the relative regulatory activity of node *i*, and represents an extension of the Linear, Latent, Causal (LLC) model [11]. This latent variable represents the effects of the phosphorylation state, but could in principle represent any post translational modification. Since phosphorylation can either activate or deactivate a regulator, it can be influenced by either kinases or phosphatases.

The edge value, *w*_*i j*_, defines the influence from node *j* to node *i* in a directed graph that may have cycles. As in the previous work, self-loops are avoided by enforcing *w*_*ii*_ = 0. *a*_*j*_(*t*) is a function of *x*_*j*_(*t*) and *y*_*j*_(*t*), which has to be defined in a meaningful way to combine the node concentration and activity attributes. 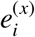 and 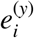 captures any latent concentration and activity contributions not explicitly mediated by the nodes in the network. In this study, *a*_*j*_(*t*) = *x*_*j*_(*t*) + *y*_*j*_(*t*) (see Derivation of Inference Model in S1 Text), and the equilibrium equations simplify to the formulation in LLC if *a*_*j*_(*t*) = *x*_*j*_(*t*).

Eqs (1a-1b) equivalently expressed in vector notation:

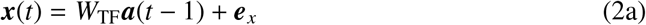

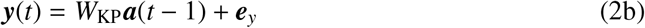

*W*_TF_ = (*w*_TF,*i j*_) and *W*_KP_ = (*w*_KP,*i j*_) are adjacency matrices containing TF edges and KP edges. Since TF ∩ KP = ∅, then *W*_TF_ and *W*_KP_ can be formulated as a single weight matrix *W* with *W*_TF_ = *WI*_TF_ and *W*_KP_ = (*I*_TF_ + *I*_KP_)*WI*_KP_, where *I*_TF_ and *I*_KP_ are diagonal matrices with ones at indexes indicating nodes in TF and KP, and zero otherwise. It has been implemented as *W*_TF_ = *W* ⊙ *M*_TF_ and *W*_KP_ = *W* ⊙ *M*_KP_, where *M*_TF_ and *M*_KP_ are indicator matrices, and ⊙ is entry-wise multiplication.

Eq (2) simplifies to the following equilibrium equations:

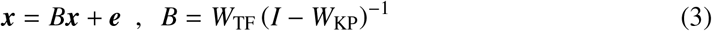

### Intervention experiments

Intervention experiments are gene perturbations, where nodes in the model (representing genes and their gene products) are knocked out or over-expressed by changing their concentration parameter. In an experiment *k*, one or more nodes in 𝒥_*k*_ (though typically one) are intervened on by setting

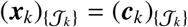

where ***c***_*k*_ is a constant intervention, and this particular subscript notation indicates subsetting of perturbed nodes (only). The combined expression for both intervened and passively observed nodes becomes:

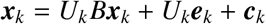

***x***_*k*_ are node values for experiment *k* which is defined by a set 𝒥_*k*_ containing indexes of intervened nodes. Multiple samples can be collected of each experiment *k*, however data is often limited to a single sample. A knockout is not affected by transcription regulation so edges in *W*_TF_ onto nodes in 𝒥_*k*_ are removed by *U*_*k*_ = (*u*_*kij*_), which is a diagonal matrix with ones indicating passively observed nodes, and zeros indicating intervened nodes (*u*_*kii*_ = 0 for *i* ∈ 𝒥_*k*_). The intervention term, ***c***_*k*_, contains zeros except for 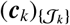 which are set to the log fold-change values measured in perturbation data. Again, ***e***_*k*_ represents noise and other latent effects.

### Loss functions

*W* can be inferred by minimizing ***e***_*k*_ for all experiments simultaneously with *L*_2_ regularization.

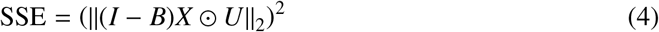

where column *k* in *X* is ***x***_*k*_, column *k* in *U* is the diagonal of *U*_*k*_, and the norm is entrywise. Minimizing SSE without regularization will result in a noisy solution. It is often corrected with an *L*_1_ penalty on the trainable weights, which are the edges in *W*. Regularization was performed on *B* instead of *W*, as regularization of *W* caused unbalanced inference. *B* was defined by the following:

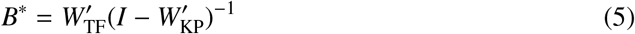

where 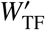 and 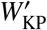 hold absolute elements of *W*_TF_ and *W*_KP_. The solution was then formulated as:

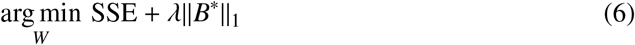

where the norm is entry-wise. All results were found using AdamW gradient descent [14] and regularization hyperparameter *λ* = 0.1.

### Restricting the solution space

The solution space can be reduced with information about the proteins. *M*_TF_ was used to disallow certain TF edges, those which lacked evidence, by setting the appropriate elements of *M*_TF_ to zero. The same was applied for *M*_KP_, where rows were set to zero for regulators which did not have evidence for at least one phosphorylation site.

If the mode of regulation is known for TF-target gene pairs, then the degrees of freedom for the solution can be reduced using 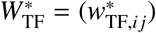 instead of *W*_TF_.

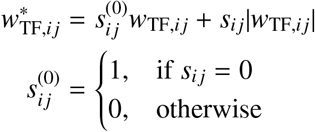

TF *j* can be an activator or repressor of gene *i*, represented with *s*_*i j*_ = 1 or *s*_*i j*_ = −1. If no prior information exists about the mode of regulation then *s*_*i j*_ = 0. The equivalent vector notation is 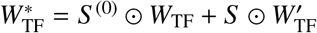, where 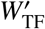 holds absolute elements from *W*_TF_.

### GeneNetWeaverPhos main equations

Data for benchmarking network inference is simulated with differential equations describing concentrations of mRNA *r*_*i*_, protein *p*_*i*_ and activated protein *ψ*_*i*_ in a cell.

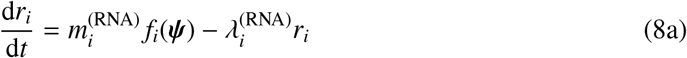

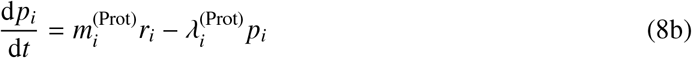

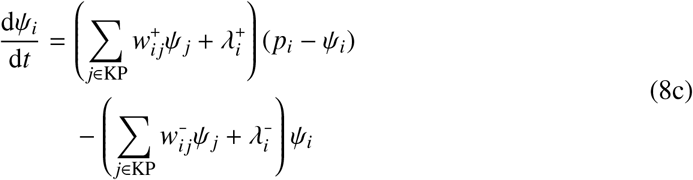

*f*_*i*_ Eq (1) in S1 Text is a nonlinear function modelling transcription regulation taking into account TF binding cooperation and competition. It holds *fi*(***ψ***) ∈ [0, 1]. 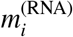 and 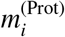 are maximum transcription and translation rates. 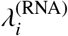 and 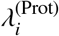 are decay rates for mRNA and protein. 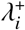 and 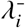 are the rate of passive activation and deactivation of protein *i*, i.e. not mediated by a specific regulator. 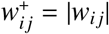 if *w*_*i j*_ > 0, otherwise 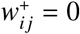. Likewise, 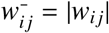 if *w*_*i j*_ < 0, otherwise 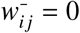.

### Modeling of transcription regulation

For the simulations performed by GeneNetWeaverPhos, the proportion of maximum transcription is used to model the regulation of a gene. For gene *i*, the function *f*_*i*_(*ψ*) uses the amount of activated regulators to estimate this proportion given the regulatory inputs to gene *i* defined in the network. The way in which information from multiple regulators was integrated is described in detail in Modeling of transcription regulation in S1 Text. Regulator concentrations for each regulatory module were combined using a generalization of the Hill equation. In the special case of a module with a single regulator, the expression for *µ*_*m*_ from Eq (1) in S1 Text simplifies to a standard Hill equation for either an activator Eq (9a) or repressor Eq (9b).

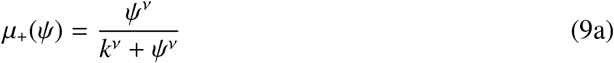

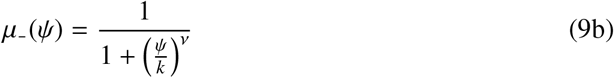

Here, *ψ* is the concentration of the transcriptional regulator which is able to bind the DNA, *k* is a dissociation constant, and *ν* is a parameter that shapes how binding sites respond to regulator saturation.

### Network construction for simulation

Five adjacency matrices from DREAM4, were each used 5 times to create 25 random adjacency matrices each with 100 nodes (see Generation of random adjacency matrices given to GeneNetWeaverPhos in S1 Text). Fully defined networks were then randomly generated with GeneNetWeaverPhos, which could subsequently be used to generate (simulated) log fold-change values. In the random networks, protein kinases and phosphatases were encoded as KPs which can both regulate positively and negatively.

### Yeast data

Many types of data were collected for inferring a regulatory network for yeast and for evaluating the performance of said inference, in an effort to validate PhosTF.

### Perturbation data

Curated perturbation data primarily consisted of gene knockouts and overexpression experiments, where (in most cases) a single gene was deleted or over-expressed (Table 2). “Tech.” refers to the technology or type of experiment performed to obtain the relative expression data, either DNA microarray (“DNA-MA”), or Sequential Window Acquisition of All Theoretical Mass Spectra (“SWATH-MS”). All measurements were log_2_ fold-change mRNA expression levels for a mutant relative to wildtype, except for SWATH-MS data which were protein measurements instead of mRNA. For the data originally published by [18], values from the reanalysis by [19] were used. “Genes” is the number of measured genes for each experiment, and “Exp.” is the number of perturbation experiments characterized. The number of different mutated genes is given in parenthesis if different from the number of experiments (due to replicates). A total of 6395 different genes were measured over the 1306 experiments. Of these, 173 different KPs were genetically manipulated (knocked out or over expressed) in 828 experiments, and 272 different TFs were similarly perturbed in 478 experiments.

**Table 2.** Yeast Perturbation Data Resources.

| Resource | Source | Tech. | Genes | Exp. |
| --- | --- | --- | --- | --- |
| KP KO | Holstege *et al.* [15] | DNA-MA | 6109 | 163 |
| V KO | Holstege *et al.* [16] | DNA-MA | 6170 | 1484 |
| KP KO | Zelezniak *et al.* [9] | SWATH-MS | 726 | 352 (97) |
| KP KO | Fiedler *et al.* [17] | DNA-MA | 6184 | 2 |
| KP KO | Hu *et al.* [18] | DNA-MA | 6253 | 5 |
| TF KO | Hu *et al.* [18] | DNA-MA | 6253 | 264 |
| TF KO | Chua *et al.* [4] | DNA-MA | 6222 | 102 |
| TF OE | Chua *et al.* [4] | DNA-MA | 6222 | 110 |

### Edge data

*d*(TF, *V*) edge data is used for *M*_*T*_ (see Equilibrium Equation in Methods), where *V* = KP ∪ TF ∪ *O*, and *O* is the set of passively observed nodes, i.e. genes with no regulatory role, with one or more TFs regulating them. *d*(KP, KPTF) edge data is used for evaluating inference performance.

“Value” displays the type of measurement if measurements were provided for the edges in the data set. Merged edge data was filtered by matching source and target nodes against mutated and measured genes from the perturbation data. “Entries” shows the number of measurements, and “Edges” is the number of edges after filtering by the sets KP, TF, and *V* (see Node Sets in Network Construction). Predictions scores were collected from NetPhorest using the provided reference yeast genome. The edge value “binding” refers to published binding evidence and “expression” refers to edges with evidence of expression regulation, where each edge only has evidence for positive or negative regulation. “Ambiguous expression” refers to edges with evidence for both positive and negative regulation. “Score” and “scores” refer to single and multiple separate arbitrary scores for each edge measuring different types of interactions, notably a score for post-translational modification. Undirected interactions allowed for a potential *d*(TF, *V*) edges in either direction.

The resulting *d*(KP, KPTF) set contained physical interaction data for 4521 of the 85371 potential edges (5.3%). Similarly, there was data for 1258143 out of 1467081 potential *d*(TF, *V*) edges (86%). Reported *p*-values from binding evidence were combined for each TF edge with Fisher’s method. If a *p*-value threshold was used but individual edge *p*-values were missing, then the *p*-value threshold was used for this set of edges (e.g. Balaji *et al*.). Similarly, YEASTRACT binding data had a conservative *p* < 0.05 restriction enforced. A False Discovery Rate threshold *q* < 0.2 was used to filter the edges for combined significance resulting in 21895 edges (∼95 per TF).

BioGRID contained data set for 40000 phosphorylation sites in 3918 proteins and another table with 111 kinases and 35 phosphatases mapped to 7561 of the sites. The BioGRID edge set was constructed through the mapping between the two tables.

### GO data

Gene Ontology data was used for categorizing the *d*(TF, *V*) mode of regulation, as well as assisting edge data in assigning proteins to sets KP, TF, and *O* (Table 4). GO Pathway terms were curated for evaluation purposes. All GO resources were from AmiGO2 version 2020-01-01 [31] except for pathway resources retrieved from SGD [32]. The “Resource” column describes the interpretation of each GO term, and “Class” shows the filtered “GO class (direct)”. In the case of Biological Process GO terms, 100 different terms were possible to test. All AmiGO2 annotation queries were filtered by organism “Saccharomyces cerevisiae S288C”. “Entries” shows the number of entries for each query and “Proteins” shows the number of proteins with at least one entry.

**Table 3.**
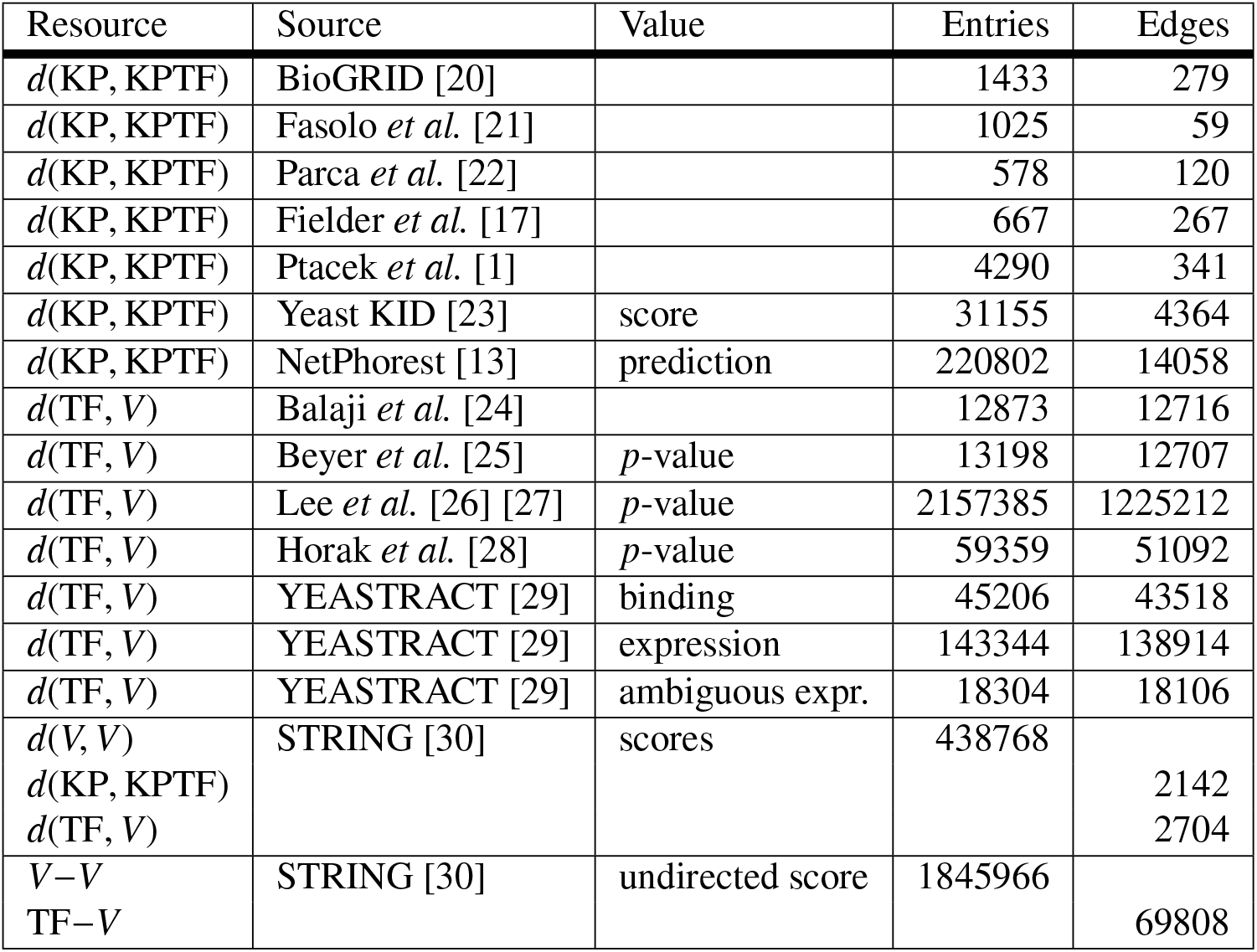
Yeast Edge Data Resources.

**Table 4.**
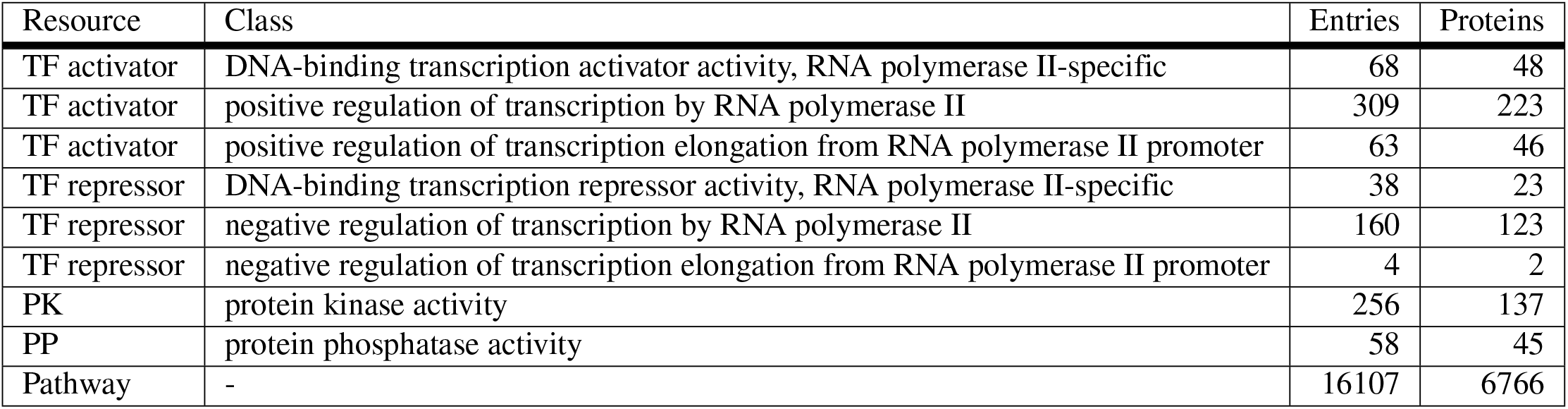
Gene Ontology Annotation Resources.

| Resource | Class | Entries | Proteins |
| --- | --- | --- | --- |
| TF activator | DNA-binding transcription activator activity, RNA polymerase II-specific | 68 | 48 |
| TF activator | positive regulation of transcription by RNA polymerase II | 309 | 223 |
| TF activator | positive regulation of transcription elongation from RNA polymerase II promoter | 63 | 46 |
| TF repressor | DNA-binding transcription repressor activity, RNA polymerase II-specific | 38 | 23 |
| TF repressor | negative regulation of transcription by RNA polymerase II | 160 | 123 |
| TF repressor | negative regulation of transcription elongation from RNA polymerase II promoter | 4 | 2 |
| PK | protein kinase activity | 256 | 137 |
| PP | protein phosphatase activity | 58 | 45 |
| Pathway | - | 16107 | 6766 |

Protein modes of regulation were classified according to curated GO evidence. GO evidence based on computational predictions alone was not considered in assigning regulatory modes. Low-throughput and direct experimental evidence was trusted over high-throughput and indirect evidence. This curation resulted in 191 TF activators, 65 TF repressors, 146 protein kinases, and 51 protein phosphatases.

### Yeast regulatory network construction

The data was processed to create an initial genome-scale regulatory network that was used for KP regulation inference. Subsections are ordered chronologically.

### Definition of node sets

The KP set was curated from the source nodes in KP interaction data as well as the mutated genes in KP perturbation data (mRNA expression profiles). The TF set was curated from the source nodes in TF interaction data filtered for target nodes in *V*, where one or more mRNA expression values were observed in perturbation data. *O* is the set of non-regulatory genes with at least one TF input (the subset of *V* not in KP or TF). Sizes of the distinct gene sets were |KP| = 199, |TF| = 231, and |*O*| = 5922.

The 40000 phosphorylation sites in BioGRID are found in 174 of the KPs and 173 of the TFs. Only these were considered valid KP substrates with potential regulatory effects, which was enforced in *M*_KP_ (rows are zero for invalid substrates).

### Node values

Expression values were averaged across replicated perturbation experiments ignoring possible differences in experimental conditions. This resulted in matrix *X* in Eq (4), while *U* was represented the mapping between manipulated and measured genes.

Perturbation effects were enhanced (see Enhancing relative expression of genetically perturbed genes in S1 Text). KOs genes were adjusted by −4, which results in an average KO gene level ∼100 times less than wildtype, and similar to values stable in simulation. OE genes were adjusted by +1, which results in an average expression ∼4 times wildtype levels.

### Initial d(TF,V) weights

TF edge weights *w*_*i j*_ represent the relationship between the log fold-change value of a source node *v*_*j*_ ∈ TF and target node *v*_*i*_ ∈ *V. w*_*i j*_ can be inferred from log fold-change values, but it is assumed that the physical binding evidence can adequately categorize TF-DNA interaction as present or absent.

TFs were categorized as either activators or repressors based on available data. The order of priority was: GO evidence, YEASTRACT and STRING combined with edge *p*-values, and lastly perturbation data. YEASTRACT and STRING described the mode of regulation for individual interactions. The *p*-values for either activating or repressing interactions were combined using Fisher’s method and compared for significance. Remaining unclassified TFs were categorized based on the average log fold-change of their targets in experiments where the TF was deleted, and if no such experiment existed, the classification was based on the sign of correlation between logFC values of the TF and its targets.

The mode of regulation for each edge was either assigned from the above listed sources, or inferred from the mode of regulation assigned to the TF. Edge weights were initialized as −1 or +1 depending on mode of regulation. All other weights were set to zero and treated as invariant during training.

### Initial d(KP,KPTF) weights

The variable (trainable) weights *w*_*i j*_ for *v*_*i*_ ∈ KPTF and *v*_*j*_ ∈ KP were initialized from a normal distribution with a small variance *σ*^2^ = 10^−4^. Initial *w*_*i j*_ for *v*_*i*_ ∈ KP and *v*_*j*_ ∈ KP were sampled randomly, however *w*_*i j*_ for *v*_*i*_ ∈ TF were informed by Wilcoxon rank tests on KP perturbation data. Absolute log fold-change values for each KP with knockout data were compared for each TF with a one-sided Wilcoxon rank test. The tests compares absolute measurements from two groups of genes; the TF regulon and remaining genes. Significant *p*-values from these tests indicate which KPs have influence on TF regulons. Instead of random sampling from a normal distribution, values were selected from a normal distribution for the quantile of the *p*-value. As a result, smaller *p*-values corresponds to larger |*w*_*i j*_|.

sgn(*w*_*i j*_) for KP *j* on TF *i* were initialized from equivalent two-sided Wilcoxon tests.

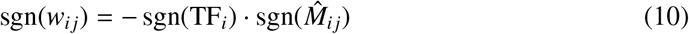

sgn(TF_*i*_) is the regulation mode of TF *i* curated from literature. 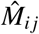 is the estimated median of difference between the two groups.

### Parameter estimation

Parameters were inferred with PhosTF for simulated data and yeast data alike. Initial states are described in Network Construction for Simulation and Network Yeast Regulatory Network Construction. Edge weights (*w*_*i j*_ ∈ ℝ) were trained on simulated data from each 100-node network by minimizing Eq (6) for 15000 epochs. In the case of training on the yeast data only 50 epochs of gradient descent were performed. Edge presence was scored as |*w*_*i j*_| and the sign was used for interpreting the mode of regulation.

### Evaluation of performance

#### Simulated regulatory networks

Performance for each inference was based on different edge weight thresholds, *θ*, where each Boolean classification generated a set of predicted present and absent edges. Prediction of an edge was either assessed as *w*_*i j*_ > *θ* for *d*(Sources, Targets, +), *w*_*i j*_ < −*θ* for *d*(Sources, Targets, −), or |*w*_*i j*_| > *θ* for *d*(Sources, Targets). When compared to the ‘true’ network edges (activating, repressing or absent), edge counts for true and false positives, and true and false negatives could be compiled. Comparing true positive rates to true negative rates, in an ROC analysis, allowed for the estimation of an area under the curve (AUC) as a measure of prediction performance.

#### Yeast regulatory networks

*d*(TF, *V*) edges were constructed from binding data, so performance was not evaluated on the inferred edge weights for these edges. The evaluation of performance on *d*(KP, KPTF) was performed using direct protein phosphorylation data since KP edges were only inferred from perturbation data, as well as indirectly implicated through the restrictions applied to *d*(TF, *V*) edges. Performance could furthermore be assessed using GO process terms since such data was also not used in the inference process.

The evaluation data set was the union of *d*(KP, KPTF) edge data sets excluding NetPhorest, and filtered for substrates with a recorded phosphorylation site (see Node Sets in Network Construction). From STRING, validation interactions were only included where evidence that a kinase phosphorylated a target protein with a protein modification (PTMod) score > 250 were included. Other than PTMod, all other lines of evidence from STRING were ignored. YeastKID was filtered with threshold score > 4.52, corresponding to *p* < 0.05, which added 412 interactions to the validation set.

Top scoring pathways were collected using thresholds of ≥ 1 to ≥ 6 shared GO terms between KP and TF from GO slim biological process terms (“Pathway” in Table 4), and are shown in Fig 4B. The 4 top *d*(KP, TF) edges (by *w*_*i j*_) were identified for each threshold: top two edges with highest and lowest edge weights (largest absolute edge weights for positive and negative regulation). The set of edges found for a particular shared GO term threshold often overlapped with those found for the other thresholds, resulting in the 16 shown.

## Discussion

This study presents a novel regulatory inference method PhosTF as well as an extension of the GeneNetWeaver simulation tool, GeneNetWeaverPhos, which includes protein kinase and phosphatase regulation of transcription factors. In addition to gaining a systems-wide understanding of how transcription factor activity is modulated by kinases and phosphatases, inferred regulatory networks can be used to predict the effects of gene mutation, or the over- or underexpression of regulators. Given further validation, such regulatory pathways can be engineered for improved control of gene expression in bioprocesses.

### PhosTF performance is state-of-the-art despite additional complexity

The prediction performance of PhosTF on simulated 100-node networks was either comparable to or better than the performance from simpler simulation models applied in the DREAM challenges. Since additional types of regulatory interactions were predicted by PhosTF, increasing the difficulty of inference, a direct performance comparison favors results from DREAM4, especially the best approaches tested in the competition. Therefore, a similar performance to the best of DREAM4 should be viewed as advancing the state-of-the-art for this more challenging inference problem.

The addition of phosphorylation and dephosphorylation activities represent hidden regulation steps and give an increased level of inference complexity. For this reason, it was surprising to find that the best prediction performance was observed for *d*(KP, TF) edges. The prediction performance of *d*(TF, *V*) was lower but still comparable to the best observed in the DREAM challenge. Since TF interactions directly regulate gene expression levels, it was expected that they would be easier to infer.

Inference on yeast was significantly harder to perform and evaluate. Evaluation was made harder by the small size of the evaluation set compared to the set of potential edges, as well as the lack of a proper negative set. With so few known examples, it cannot be assumed that the majority of novel predictions are false positives. And although there may be some false interactions in the validation set, the biggest challenge was clearly the lack of existing knowledge and inability to assess true negative rates. This means that prediction specificities cannot be realistically estimated. Sensitivity estimates were 0.40 for *d*(KP, TF) and 0.28 for *d*(KP, KP) predictions meaning that roughly 30-40% of what was known was predictable from the data and this approach, see Fig 3. Nevertheless, the enrichment of known interactions in the prediction set was highly significant. We conclude that PhosTF performance is significantly better than random predictions, and outperformed the existing kinase specificity based predictions of NetPhorest. Of the 8610 new *d*(KP, TF) and 6299 new *d*(KP, KP) predicted regulatory interactions, 30-40% would be expected to be real either through direct activity or though other indirect regulatory effects.

### Phosphorylation more often negatively regulates transcription factors

A large majority of protein kinase regulatory influences on transcription factors (71%) appeared to be negatively regulating (deactivating) these transcription factors through either direct of indirect means. This bias was even stronger for the 166 predicted *d*(KP, TF) edges that were already known, and believed to be direct interactions, where 85% were estimated to have a deactivating effects. Despite this surprising predominance of negative regulation by KPs (greater than 2-to-1), the higher than expected prediction of de-repression pathways suggests selection of transcriptional activation by protein kinases through the negative regulation of repressors.

It is tempting to assume this bias was due to systematic aspects of the gene expression data for the various knock-outs used for inference, i.e. where differentially expressed genes were non-specifically affected by gene loss. Interestingly, non-specific deletion effects would only be expected to increase the two cases where KP regulation of TFs was positive (Pos.-Pos. and Pos.-Neg. Fig 1) as these are the two type of pathways where the knockouts of the KP and the TF induce the same type of KO-affect on the target genes. In contrast, for both Neg.-Pos. and Neg.-Neg. pathways, negative regulation of transcription factors requires that the transcriptionally regulated target genes change the sign of their differential expression. Thus, non-specific or consistent KO effects would be expected to implicate positive KP regulation half of the time. As a further check, the signs of differential expression for all perturbation measurements were compared between KP and TF perturbations for each predicted *d*(KP, TF) edge. The comparisons of signs across knockout profiles did not reveal systematic anti-correlation of transcriptional responses suggesting such biases were not expected by random chance.

One possible explanation for the over representation of negative KP regulation could relate to the nucleocytoplasmic trafficking of TFs as a function of their phosphorylation state. Protein phosphorylation is known to regulate trafficking of proteins in and out of the nucleus, and is particularly relevant for TFs as this is where their primary mode of action takes place. The trafficking hypothesis would imply that phosphorylation more often facilitates retention of TFs in the cytoplasm, effectively deactivating them. This idea is not well supported by our current knowledge of nucleocytoplasmic trafficking, where are as many or more examples describe phosphorylation promoting nuclear import than cytoplasmic retention [33]. Despite anecdotal evidence to the contrary, many examples are known where phosphorylation of TFs facilitates nuclear export, e.g. Pho4p, Mig1p and Crz1p [34] suggesting that this is still a plausible explanation.

### Inference limitations

Despite the relatively large number of regulatory interactions that could be predicted from our approach, a number of possible inference challenges were identified. One such consideration that can influence the ability to infer regulation is the concept of silent regulation. Silent regulation can occur if activation or deactivation of a protein is made undetectable by the deactivation of a node downstream in a cascade. The edge weights are initially set with random values, and if they are too small (weak), a signal along a cascade can disappear, and if they are too large (strong) the nondimensionalized quantities can be forced to their bounds [0, 1]. Naturally, deactivation of a node with activity 0 will not be detectable, and vice-versa for activation of a node with the maximum activity 1. Some silent regulation is expected to be present in the biological data as well. For example a knockout can affect multiple regulatory pathways, which competitively regulate a shared target. The overall effects can cancel each other out, leaving both pathways unobservable. Some steps were taken to avoid silent regulation in the simulations, e.g. setting the magnitudes lower for edges sharing the same target (see Generation of random adjacency matrices given to GeneNetWeaverPhos in S1 Text). Silent regulation is not always avoided, which means some regulation cannot be inferred.

In addition, some regulatory effects can be silent due to compensatory pathways. Compensating signal transduction cascades are difficult to infer from perturbation data if only a single gene has been deleted from either cascade. The other cascade can compensate for the missing regulation, resulting in minor or indistinguishable changes in the expression levels of regulated targets. This limitation can be overcome with multiple knockouts in the same experiment, however most of the collected data was from single gene knockouts.

### Information attainable from genetic perturbation

The experimental conditions used for assessing the effects of genetic perturbations also has an impact on inference. This is because certain regulatory signal cascades will only be activated under specific environmental conditions or stresses, and as such will be difficult to detect from mutants measured under nominal conditions. Measurements from different conditions have been averaged, but many other approaches exist, such as averaging separately for each condition, or keeping all original experiments. Many genes are essential for survival, requiring knockdowns or conditional knockouts, further altering the experimental conditions. However, the data used here were from different labs, and in some cases included only smaller subsets of genes, as opposed to genome-wide profiles, which would result in a high number of missing values if not averaged with other replicates.

PhosTF inference relies on information from the genetic perturbation data, which is only indirect evidence of the protein regulatory interactions that are inferred. Ambiguous solutions can arise even in simple network models using simulated perturbations. Fig 2 illustrates a case where two kinases, KP_1_ and KP_2_, have identical influence (Fig 2B), which makes it impossible to infer if one regulates the other or if both regulate TF_1_, only from observing perturbation data. When multiple solutions exists the most simple is inferred, where simplicity is rewarded through regularization. Usually the *L*_1_ norm is applied to induce sparsity among trainable weights, however this assumes each weight can be treated equally. In LLC, influence from a node propagates through the network as a series of multiplications of edge values *w*_*i j*_, see Eq (1b). It usually holds |*w*_*i j*_| < 1, which helps convergence but also causes the decay of a signal along a cascade. As a result, sparsity may suffer when multiple shorter cascades are preferred over fewer, but longer, cascades.

To induce sparsity, edges can instead be penalized based on their final influence on gene expression rather than on their influence on the next node in a cascade, which is accomplished by regularizing *B*^∗^ instead of *W* in Eq (6). From testing on small simulated networks, it was found that PhosTF performs much better when the regularization was applied to *B*^∗^ instead of *W*, where regularization is typically performed. This is likely because *L*_1_ regularization of *W* unevenly penalizes *d*(KP, KP) edges compared to *d*(KP, TF) and *d*(TF, KPTF) edges (KPTF = KP ∪ TF). This improvement alone is likely why our prediction performance compares favorably to previous DREAM winners despite the additional challenges imposed by kinase regulation.

### Further work

Although edges were inferred based on the score |*w*_*i j*_|, this score does not readily describe a probability for the presence of a protein-protein interaction. A weak interaction could be accurately inferred but discarded for its low |*w*_*i j*_|. Alternative scoring methods can be considered to estimate the probability of edge presence, for instance running PhosTF multiple times and taking the variation of *w*_*i j*_ into account.

PhosTF can be combined with other methods applying evolutionary information, where knowledge of one species is transferred to another related, but less studied species [35]. A cost function for evolutionary distance can be incorporated into an ensemble method. Likewise, other data types could be considered for training, e.g. other functional knowledge or using the evaluation data set, which could improve the inference performance instead of simply evaluating it.

The simulation method GeneNetWeaverPhos has potential future use in inference approaches. Given enough computational resources, GeneNetWeaverPhos could be iteratively run with variations to the network structure, to minimize the difference between simulated and experimental gene expression levels. This can be accomplished with a Markov chain Monte Carlo approach such as the Metropolis-Hastings algorithm.

## Supporting information

**S1 Text. Method details**. GeneNetWeaverPhos simulation details and derivation of inference model.

## Data and code availability

All source code, data and results are available online at github.com/degnbol/PhosTF.

## Author contributions

CTW defined the problem and conceived the design of the initial study. CDM implemented the code for the inference and performed all analyses and compiled the data sets. CDM wrote the initial version of the manuscript. CTW and JH aided with the interpretation of the results and contributed to the manuscript revisions.

